# Multi-synaptic boutons are a feature of CA1 hippocampal connections that may underlie network synchrony

**DOI:** 10.1101/2022.05.30.493836

**Authors:** Mark Rigby, Federico Grillo, Benjamin Compans, Guilherme Neves, Julia Gallinaro, Sophie Nashashibi, Gema Vizcay-Barrena, Florian Levet, Jean-Baptiste Sibarita, Angus Kirkland, Roland A. Fleck, Claudia Clopath, Juan Burrone

**Affiliations:** MRC Centre for Neurodevelopmental Disorders, Institute of Psychiatry, Psychology and Neuroscience, King’s College London, London, United Kingdom SE1 1UL; Centre for Developmental Neurobiology, Institute of Psychiatry, Psychology and Neuroscience, King’s College London, London, United Kingdom SE1 1UL; The Rosalind Franklin Institute, Harwell Campus, Didcot, OX11 0FA, United Kingdom; Bioengineering Department, Imperial College London, London, United Kingdom; Centre for Ultrastructural Imaging (CUI), Kings College London, New Hunts House, Guys Hospital Campus, London SE1 1UL, United Kingdom; Univ. Bordeaux, CNRS, Interdisciplinary Institute for Neuroscience, IINS, UMR 5297, F-33000 Bordeaux, France; Univ. Bordeaux, CNRS, INSERM, Bordeaux Imaging Center, BIC, UAR3420, US 4, F-33000 Bordeaux, France

**Author notes:** Equal contribution (Mark Rigby, Federico Grillo).

## Abstract

Excitatory synapses are typically described as single synaptic boutons (SSBs), where one presynaptic bouton contacts a single postsynaptic spine. Using serial section block face scanning electron microscopy, we found that this textbook definition of the synapse does not fully apply to the CA1 region of the hippocampus. Roughly half of all excitatory synapses in the *stratum oriens* involved multi-synaptic boutons (MSBs), where a single presynaptic bouton containing multiple active zones contacted many postsynaptic spines (from 2 to 7) on the basal dendrites of different cells. The fraction of MSBs increased during development (from P21 to P100) and decreased with distance from the soma. Curiously, synaptic properties such as active zone (AZ) or postsynaptic density (PSD) size exhibited less within-MSB variation when compared to neighbouring SSBs, features that were confirmed by super-resolution light microscopy. Computer simulations suggest that these properties favour synchronous activity in CA1 networks.

## Introduction

In mammalian brains, chemical synapses are usually thought of as bi-cellular units formed by a single presynaptic bouton contacting a single postsynaptic compartment. These connections are typically referred to as single synaptic boutons (SSBs) and are widespread throughout the brain. However, a closer look at synaptic morphologies from electron microscopy studies show that alternative arrangements also exist (Walmsley et al., 1998). They include single postsynaptic compartments that receive multiple presynaptic inputs (multi-synaptic spines or MSSs) and single presynaptic boutons that form synapses onto multiple postsynaptic sites (also known as multi-synaptic boutons or MSBs) (Peters et al., 1991). Even though these apparently rarer configurations represent an interesting alternative to the prototypical synapse, their distribution and prevalence in the brain, as well as their morphological properties and functional role in neuronal circuits, are not well established.

MSBs are a particularly curious configuration that have been intimately associated with long-term forms of structural plasticity in the brain (Bourne and Harris, 2012; Harris, 1995). For example, the number of MSBs in the cerebellar and motor cortices has been shown to increase following the emergence or modulation of complex motor skills (Federmeier et al., 2002; Jones, 1999) and can be actively modulated in the visual cortex by either sensory deprivation (Friedlander et al., 1991) or exposure to complex visual scenes (Jones et al., 1997). Similarly, associative learning paradigms (eye blink conditioning) in rabbits showed an increase in the fraction of MSBs in the *stratum radiatum* of the hippocampal CA1 subfield (Geinisman et al., 2001). However, the mechanisms responsible for the appearance of MSBs are less well understood. Early studies showing that estrogen increased the density of spines in the hippocampus, also found that it resulted in an increase in the number of MSBs, suggesting that new spines may target existing boutons to create MSBs. Indeed, work in the cortex performing correlative *in vivo* imaging of spines together with EM reconstructions showed that newly formed spines were preferentially associated with MSBs (Knott et al., 2006). In line with this, spinogenesis induced by LTP-like stimuli in hippocampal slices observed newly formed spines synapsing onto pre-existing boutons (Nagerl et al., 2007; Toni et al., 1999). In this context, the presynaptic bouton of an MSB can be seen as an anchor point for new spines to latch onto to form new synaptic connections. Functionally, however, the role played by MSBs is not well understood. In the monkey prefrontal cortex, the modulation of MSB number with estrogen was shown to correlate with behavioural measures of working memory, suggesting that MSBs may play some role in cognitive function. Together with the activity-dependent plasticity described above, MSBs appear to be an important, yet understudied neuronal structure linked to key physiological processes in the brain.

EM reconstructions have shown that although MSBs appear to be present in many brain areas, including the hippocampus and cortex, they are typically in the minority (Bourne and Harris, 2008; Kasthuri et al., 2015; Knott et al., 2006; Sorra and Harris, 1993). Here, we show that for excitatory synapses in the *stratum oriens* (SO) of the hippocampal CA1 subfield, MSBs are present in roughly equal proportion to SSBs. What is more striking is that these MSBs involve single boutons that form contacts onto multiple spines from different dendrites of different cells. By looking at EM reconstructions of two different ages (P22 and P100) we show that MSBs are not a developmental anomaly but instead represent a feature of how CA1 hippocampal neurons receive inputs. Moreover, the properties of synaptic contacts within single MSBs are less heterogeneous than that observed across neighbouring SSBs, suggesting MSBs convey similar information across its many contacts. Using mathematical models of network activity that incorporate these properties, we propose that MSBs in the hippocampus act to synchronise network activity by using single boutons to broadcast information to multiple postsynaptic partners.

## Results

We set out to look at the properties of synapses in the *stratum oriens* of the CA1 hippocampus, a region populated by basal dendrites, which receive a large fraction of the total number of inputs onto CA1 pyramidal neurons. We performed serial EM reconstructions of two regions of the *stratum oriens*, one proximal and another distal to *stratum pyramidale* (Fig. 1A) for both a young adolescent (P22) and an adult (P100) mouse brain. In each case we reconstructed dendrites and axons within 1000 μm^3^ volumes (a cube of 10×10×10 μm) allowing us to establish the properties of synapses along the distance of basal dendrites to the soma, as well as across development. Having reconstructed hundreds of synapses we noticed a salient feature in the properties of presynaptic terminals in the *stratum oriens*. Data pooled from all our reconstructions showed that nearly half (^~^45%) of the boutons in this area formed synapses onto more than one postsynaptic spine, resulting in structures known as MSBs. More intriguingly, the spines contacted by MSBs belonged to different dendrites, rather than neighbouring spines on the same dendrite (Fig. 1B). From the general topology of dendritic arbours we predicted that these dendrites likely belonged to different neurons, suggesting MSBs were transmitting signals across different cells. To confirm this, we reconstructed neurons sparsely labelled with fluorescent proteins and mapped the 3D spatial arrangement of their dendrites (Fig. S1). We found that dendrites tended to fan out from the soma along the *stratum oriens* (Fig. S1A-H), rarely coming into close contact with each other. A measure of potential contacts with dendrites from the same cell decreased with distance from the soma, becoming negligible beyond ^~^50 μm (Fig. S1I). Since the most proximal EM data set was obtained at a distance of ^~^50 μm (Fig. 1A), dendrites from the same cell were unlikely to contribute more than a single spine to an MSB. Together, our data strongly suggests that the vast majority, if not all, of the MSBs described here represent a single presynaptic bouton contacting the dendrites of multiple postsynaptic CA1 pyramidal neurons. Although a single MSB can contact up to 7 postsynaptic spines, the majority ranged from 2 to 5 contacts (Fig. 1C), with the proportion of MSBs decreasing progressively with the number of postsynaptic partners they contacted. Nevertheless, MSBs forming more than 3 contacts still represented as much as 15% of all boutons along a dendrite or axon (Fig. 1C). Morphologically, bouton volume scaled with the number of synapses formed, with boutons progressively increasing in size the more active zones they had (Fig. 1D). We conclude that MSBs are relatively large structures that are abundant in the *stratum oriens* of the hippocampus and connect multiple CA1 neurons together.

**Figure 1.**
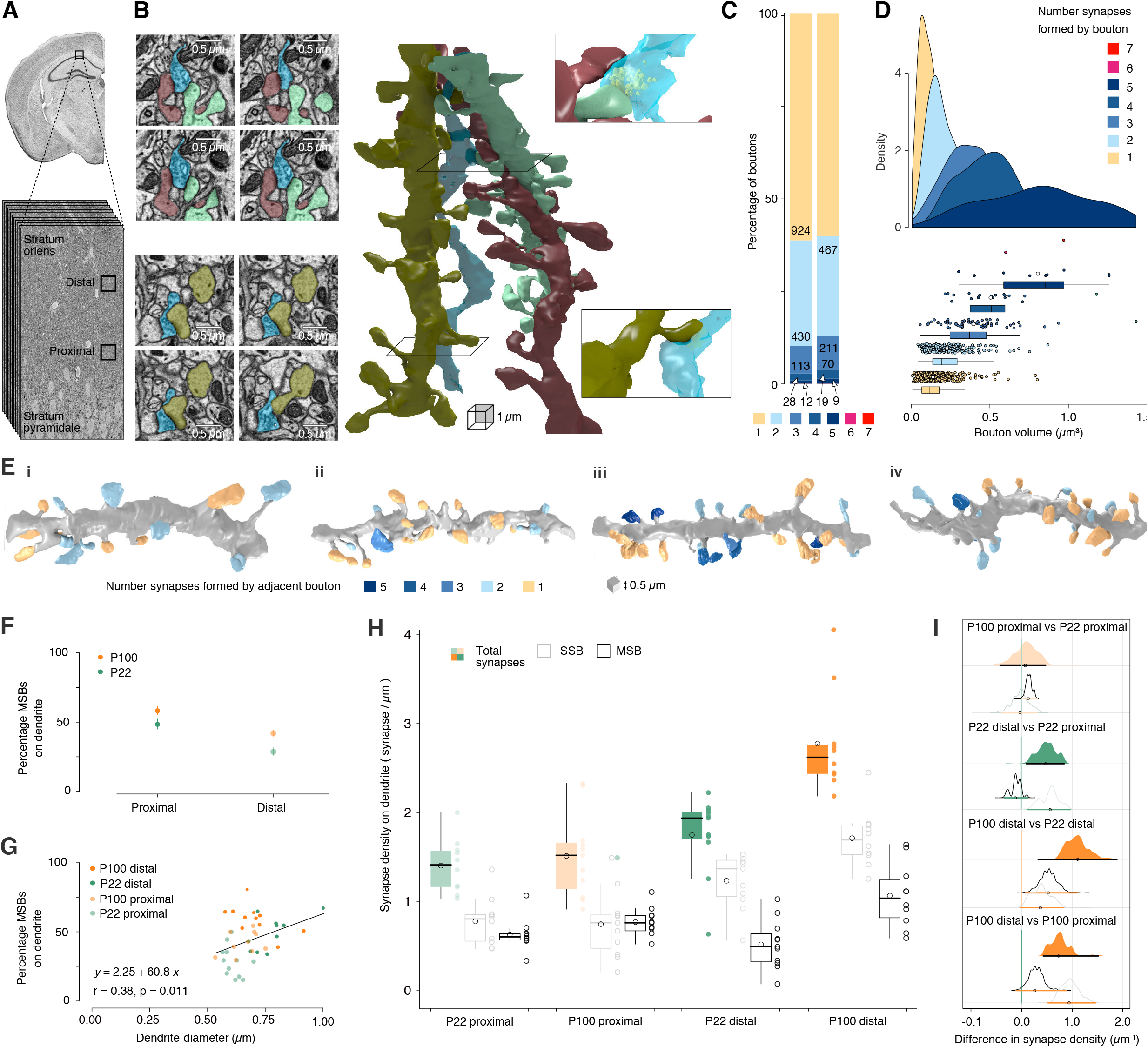
Serial block face scanning electron microscopy reveals multi synaptic boutons feature heavily in the *stratum oriens*. (**A**) Low magnification image of *stratum oriens* and stratum pyramidale of the hippocampus, where basal dendrites of CA1 pyramidal neurons reside. Black box represents the cubes of tissue extracted and imaged using serial block face scanning EM. 2 cubes, 1 proximal to the stratum pyramidale and another more distally located from the CA1 somata region were processed from a P22 mouse and also from a P100 mouse. (**B**) 3D reconstructions of 3 dendrites (dark grey, olive green & bright mint green) forming synapses onto a single axon (blue). Example images on the left and close up 3D reconstructions on the right are the corresponding slices through the z-stack. The top example shows a multi synaptic bouton forming 2 connections, whereas the bottom images and reconstruction are of a single-synaptic bouton. (**C**) Boutons from both animals were classified depending on how many synaptic contacts they formed and plotted as a percentage of the total number of boutons. The left bar refers to boutons that form onto traced dendrites, whereas the right bar refers to boutons analysed from selected axons. The colour code represents the number of synapses formed by a bouton (legend in D). (**D**) Density histogram of axonal bouton volume grouped by the number of contacts made. The greater the number of contacts formed, the greater the bouton volume (Krusal-Wallis p value: 1.82e-50, Omega squared: 0.312). (**E**) 3D reconstructions of dendrites traced in the *stratum oriens* of 4 mice; P22 proximal (i), P22 distal (ii), P100 proximal (iii), and P100 distal (iv). Spine colour indicates the number of contacts formed by the apposing bouton. Yellow refers to a spine forming onto a single synaptic bouton, blue spines represent multi synaptic boutons, the darker the shade of blue, the more synapses formed by the apposing multi synaptic bouton. (**F**) Plot of the percentage of MSBs on either proximal or distal dendrites, for both P22 (green) and P100 (orange) dendrites. (**G**) Plot of the percentage of MSBs as a function of dendrites diameter for all dendrits analysed (P22 green and P100 orange) showing a correlation between the two. (**H**) Pooled data showing the density of total synapses formed on individual dendrites (filled), as well as the density of synapses broken down into those formed by single synaptic boutons (grey outline) and multi synaptic boutons (black outline). Box-and-whisker plots indicate the median (black line), the 25th–75th percentiles (black box) and the 10th–90th percentiles (black whiskers); open circles within the box and to the right of the box represent the mean and individual values, respectively. (**I**) Age and region comparisons of synaptic densities visualised using estimation plots of effect sizes and their uncertainty using confidence intervals Medians are indicated with a circle and the 95% confidence intervals with a horizontal line. Separation between the confidence interval and 0 was considered an effect. Confidence intervals were adjusted for multiple comparisons by an extension of the Ryan–Holm stepdown Bonferroni procedure. 4 comparisons were made for total synaptic density, whereas confidence intervals were adjusted for 12 comparisons for synapse densities formed by single and multi synaptic boutons.

Previous work has shown that the properties of synaptic boutons follow distance-dependent rules along the basal dendrites of CA1 pyramidal neurons (Grillo et al., 2018). We therefore also explored the possibility that MSBs may follow a non-random spatial distribution. We found that the proportion of MSBs was highest in proximal dendrites for both P22 and P100 brains, reaching surprisingly high levels (>50%) in the proximal dendrites of the P100 brain (Fig. 1E-F). Although the proportions remain high all along basal dendrites, distal dendrites showed a decrease in the fraction of MSBs, suggesting there are distance-dependent rules that control the likelihood of finding an MSB in the *stratum oriens* (Fig. 1F). Basal dendrites display a pronounced tapering with distance along the *stratum oriens*, and previous studies have used dendrite diameter as a proxy for measuring the distance of basal dendrite from the soma (Grillo et al., 2018; Walker et al., 2017). We found that the proportion of MSBs also correlated with dendrite diameter, suggesting a graded decrease in the fraction of MSBs along dendrites, when moving away from the soma (Fig. 1G).

A more intriguing picture emerged when looking at the absolute density of synapse types along dendrites. Distal dendrites showed an increase in overall spine density when compared to proximal dendrites, which was more apparent for older (P100) neurons and was mainly explained by an increase in distal SSBs (Fig. 1H-I). In fact, there was a clear developmental increase in overall synapse density (for both SSBs and MSBs) from P22 to P100 in distal dendrites, suggesting synapse formation continues beyond the third postnatal weeks in this area of the hippocampus (Fig. 1H-I). Hierarchical clustering of dendritic domains based on SSB and MSB densities uncovered essentially 2 main groups populated by mostly proximal or distal domains, regardless of age (Fig. S2). This distinction between proximal and distal dendritic domains is in line with previous findings showing distance-dependent differences in the synaptic properties of both postsynaptic spines and presynaptic boutons (Grillo et al., 2018; Walker et al., 2017) and strengthens the idea that different dendritic domains code for different types of information. Here, it appears that although MSBs are found in roughly equal numbers at all dendritic distances, the proportion of MSB synapses is biased towards proximal domains.

What are the properties of postsynaptic spines that connect to either MSBs or SSBs? Morphological measures of dendritic spines (Fig. 2A) are intimately linked to their function: both spine volume and PSD size correlate well with postsynaptic strength and have therefore regularly been used as structural measures of spine function (Matsuzaki et al., 2001; Zito et al., 2009). We found that at P22, spines contacting MSBs and SSBs were very similar in terms of volume and PSD size, whereas at P100, MSB spines were smaller (Fig. 2B-E), in line with previous work in the *stratum radiatum* (Nicholson and Geinisman, 2009). In general, morphological measures of postsynaptic spines increased in size from P22 to P100 for both MSBs and SSBs, particularly in proximal domains (Fig. 2C and E). There was also an interesting spatial distribution of postsynaptic spine properties, where spines onto both SSBs and MSBs showed an overall decrease in volume and PSD size with distance along a dendrite (Fig. 2C and 2E), in agreement with previous observations (Grillo et al., 2018; Walker et al., 2017). Together, our data shows that spines formed onto MSBs are generally similar to SSBs at P22, but smaller at P100, and that dendritic location is an important determinant of spine size across all ages and synapse types. Synapses are highly heterogenous entities, with both pre- and postsynaptic compartments showing large variations in structure and function (Dobrunz and Stevens, 1997; Grant and Fransen, 2020). Although this heterogeneity is clearly present across both SSBs and MSBs, the variability in synaptic properties within single MSBs is less well understood. We therefore calculated the variance for AZ size, PSD size and spine volume of contacts within MSBs (for MSBs ranging from 2 to 5 contacts) and compared them to neighboring SSBs (from 2 to 5 neighbours) on a dendrite. Surprisingly, we found that all synaptic measures were more similar to each other within an MSB than across SSBs (Fig. 2F-I). To explore this further, we carried out dual-colour direct Stochastic Optical Reconstruction Microscopy (dSTORM) of two presynaptic molecules, vGlut-1 and Bassoon, to uncover the 3D nanoscale arrangement of vesicles and their AZs, respectively, and to sample the properties of MSBs across multiple brains (Fig. 3). Whereas staining for vGlut-1 showed clouds of vesicles that outlined presynaptic boutons, bassoon staining was predictably more punctate (Fig. 3A-B). Once again, we found that a large proportion of boutons (^~^ 25 %) contained multiple active zones, ranging from 2 to 7. The lower numbers MSBs identified with this method is likely due to a conservative approach to assigning AZ puncta to single vGlut clusters (Fig. S3), as well as to limitations in the resolution and imaging depth of 3D dSTORM in slices, which will strongly underestimate the number of AZs detected. In some cases, the location of AZ puncta were clearly identified at opposite ends of a bouton, suggesting that postsynaptic partners were likely to be on different spines, as observed for MSBs using SBFSEM. In line with the morphological characterization of bouton volume (Fig. 1D) and AZ size, we found that MSBs had larger vGlut cluster volumes (Fig. 3C) but no change in bassoon cluster volumes (Fig. 3D) when compared to SSBs. More importantly, we found that bassoon cluster volume was less variable within MSBs than across SSBs (Fig. 3E), in agreement with the decrease in variation measured for MSB AZ size. Together, our data suggests that MSBs transmit similar information at each of its contacts to multiple cells. Particularly, the similar AZ size is indicative of similar release probabilities, which would also imply similar short-term dynamics of neurotransmitter release.

**Figure 2.**
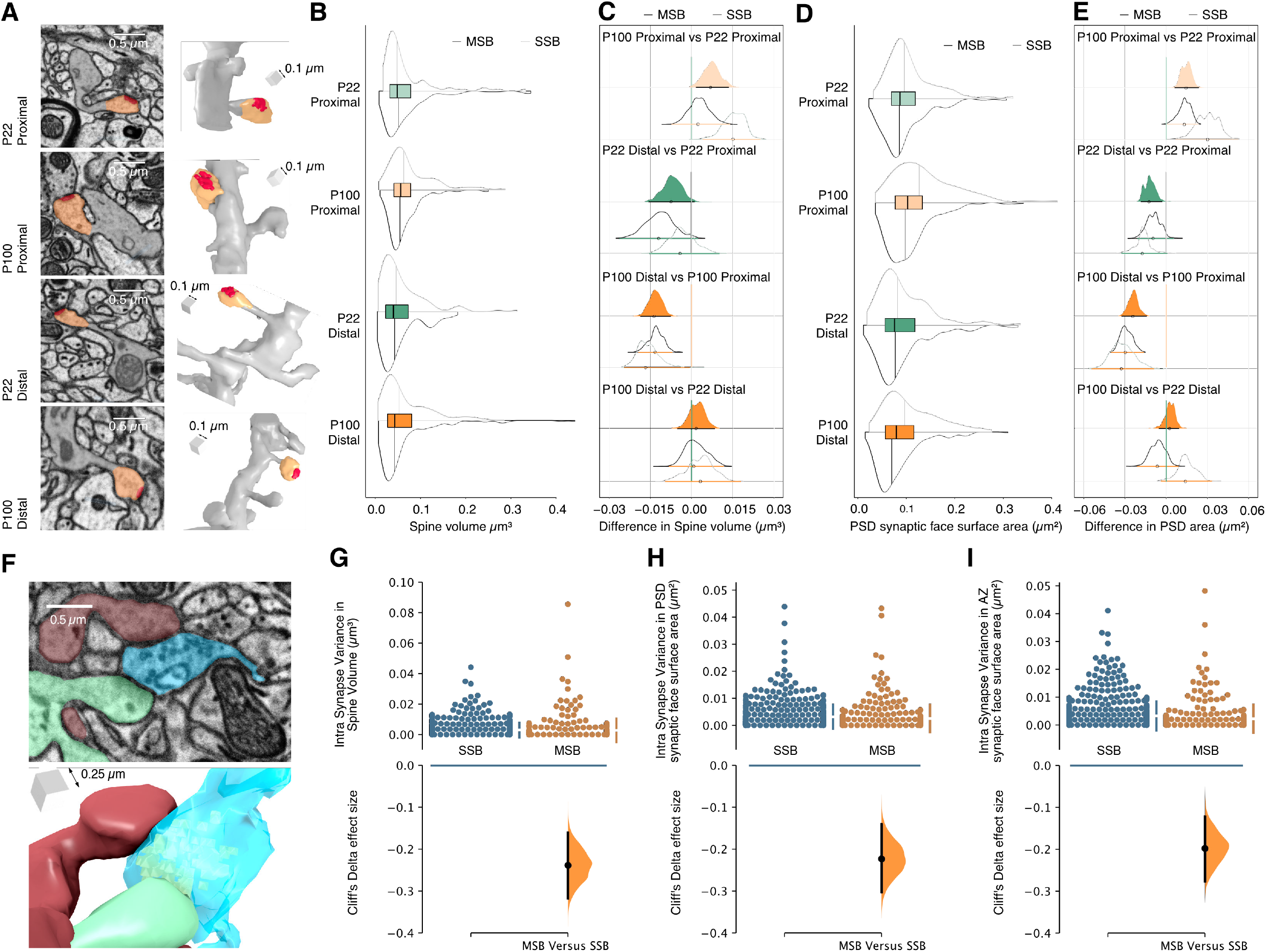
Properties of synapses formed by MSBs. (**A-E**) Synapse strength increases from distal to proximal dendrites in the adult but not juvenile mouse. (**A**) Representative electron micrographs and corresponding 3D reconstructions of individual dendritic spines taken from mice of different ages and regions in the *stratum oriens*. (**B**) Combined plot describing the distributions of spine volumes for the total population (box plot; as described above), those dendritic spines formed onto single synaptic boutons (grey split violin plot) and dendritic spines formed onto multi synaptic boutons (black split violin plot). (**C**) Estimation plot of spine volume to visualise effect sizes and their uncertainty Medians are indicated with a circle and the 95 confidence intervals with a horizontal line. Separation between the confidence interval and 0 was considered as an effect. Confidence intervals were adjusted for multiple comparisons by an extension of the Ryan–Holm step-down Bonferroni procedure. (**D**) Combined plot describing the distributions of PSD synaptic face surface areas for the total population (box plot; as described above), those dendritic spines formed onto single synaptic boutons (grey split violin plot) and dendritic spines formed onto multi synaptic boutons (black split violin plot). (**E**) Estimation plot of PSD synaptic face surface areas to visualise effect sizes and their uncertainty. Medians are indicated with a circle and the 95% confidence intervals with a horizontal line. Separation between the confidence interval and 0 was considered an effect. Confidence intervals were adjusted for multiple comparisons by an extension of the Ryan–Holm step-down Bonferroni procedure. (**F-J**) Synaptic properties exhibit greater homogeneity when from the same MSB than across individual SSBs on the same dendrite. (**F**) Representative electron micrograph and corresponding 3D reconstruction of multi synaptic bouton with 2 connecting spines. (**G**) Quantification of the variance in spine volume for neighbouring SSBs (blue) and for spines formed onto single MSBs (orange). Vertical error bars indicate the 95% CI (two-sided permutation t test). Cliff’s Delta effect size between MSBs and SSBs variance is shown in the bottom panel. (**H**) Quantification of the variance in PSD size for spines on neighbouring SSBs (blue) and for spines formed onto single MSBs (orange). Vertical error bars indicate the 95% CI (two-sided permutation t test). Cliff’s Delta effect size between MSBs and SSBs variance is shown in the bottom panel. (**I**) Quantification of the variance in AZ size for neighbouring SSBs (blue) and for AZs within a single MSB (orange). Vertical error bars indicate the 95% CI (two-sided permutation t test). Cliff’s Delta effect size between MSBs and SSBs variance is shown in the bottom panel. Data from all mice are pooled together.

**Figure 3.**
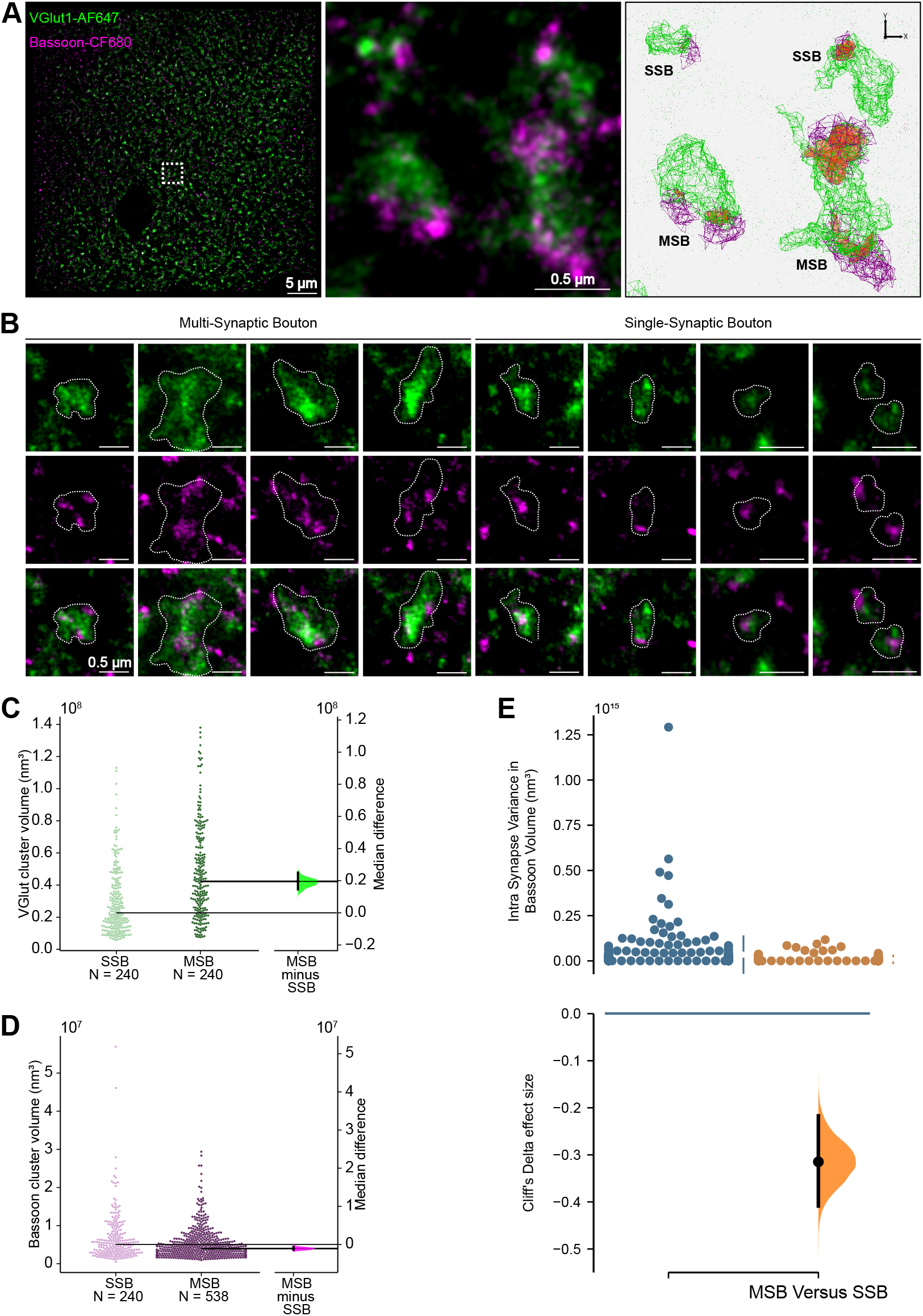
Super-resolution imaging reveals that AZ properties within a MSB are more similar than between SSBs. (**A**) 2 colour 3D-dSTORM image of VGlut (green) and Bassoon (magenta) in the *stratum oriens* of a P22 mouse brain. Middle panel shows a zoomed-in view of four VGlut boutons (green) and their corresponding Bassoon-labelled AZ puncta (magenta). Right panel shows the clusters identified by the Point Clouds Analyst (PoCA) approach used for automatic segmentation of vGlut and Bassoon, with the overlap between segmented clusters shown in orange. (**B**) Examples of Multi-Synaptic and Single-Synaptic Boutons. (**C**) Quantification of VGlut cluster volume for SSBs and MSBs. Gardner– Altman estimation plot indicate the median difference in VGlut cluster volume between MSBs and SSBs. (**D**) Quantification of Bassoon cluster volume for SSBs and MSBs. Gardner–Altman estimation plot indicate the median difference in Bassoon cluster volume between MSBs and SSBs. (**E**) Quantification of the intra-synapse variance of Bassoon cluster volume for SSBs (blue) and MSBs (orange). Vertical error bars indicate the 95% CI (two-sided permutation t test). Cliff’s Delta effect size between MSBs and SSBs variance is shown in the bottom panel.

In order to explore the functional role that MSBs could play in the hippocampus, we performed computer simulations in which CA1 neurons received synaptic input from CA3/2 neurons as well as background input. We hypothesized MSBs could influence the synchronization of CA1 pyramidal cells through three different mechanisms, namely a multiplicative effect (increased connectivity), release probability properties and short-term plasticity (Fig. 4A). We modeled the multiplicative effect by connecting each bouton from CA3/2 neurons to either a single neuron (SSB) or to multiple neurons (MSB) in CA1. To match the lower variance of AZ size within MSBs, we assumed that each bouton had a fixed release probability, drawn independently for each bouton, so that all AZs within a single MSB had an identical Pr. Furthermore, all synapses were subject to short-term plasticity, and each bouton was either facilitating or depressing, such that all AZs within a single MSB would have the same STP properties. Our results showed that, for such a configuration, the correlation between activity of neurons in CA1 (measured as the correlation coefficient *μ_cc_*) is higher in networks with MSBs, when compared to networks with SSBs only (Figure 4B). This model implicitly assumes that the short-term plasticity state of each bouton is independent of their release probability. However, Pr is known to correlate well with STP at classical synapses. We therefore matched the Pr of a bouton with an appropriate STP (high Pr boutons-depression; low Pr boutons - facilitation) and found similar results (Figure S4).

**Figure 4.**
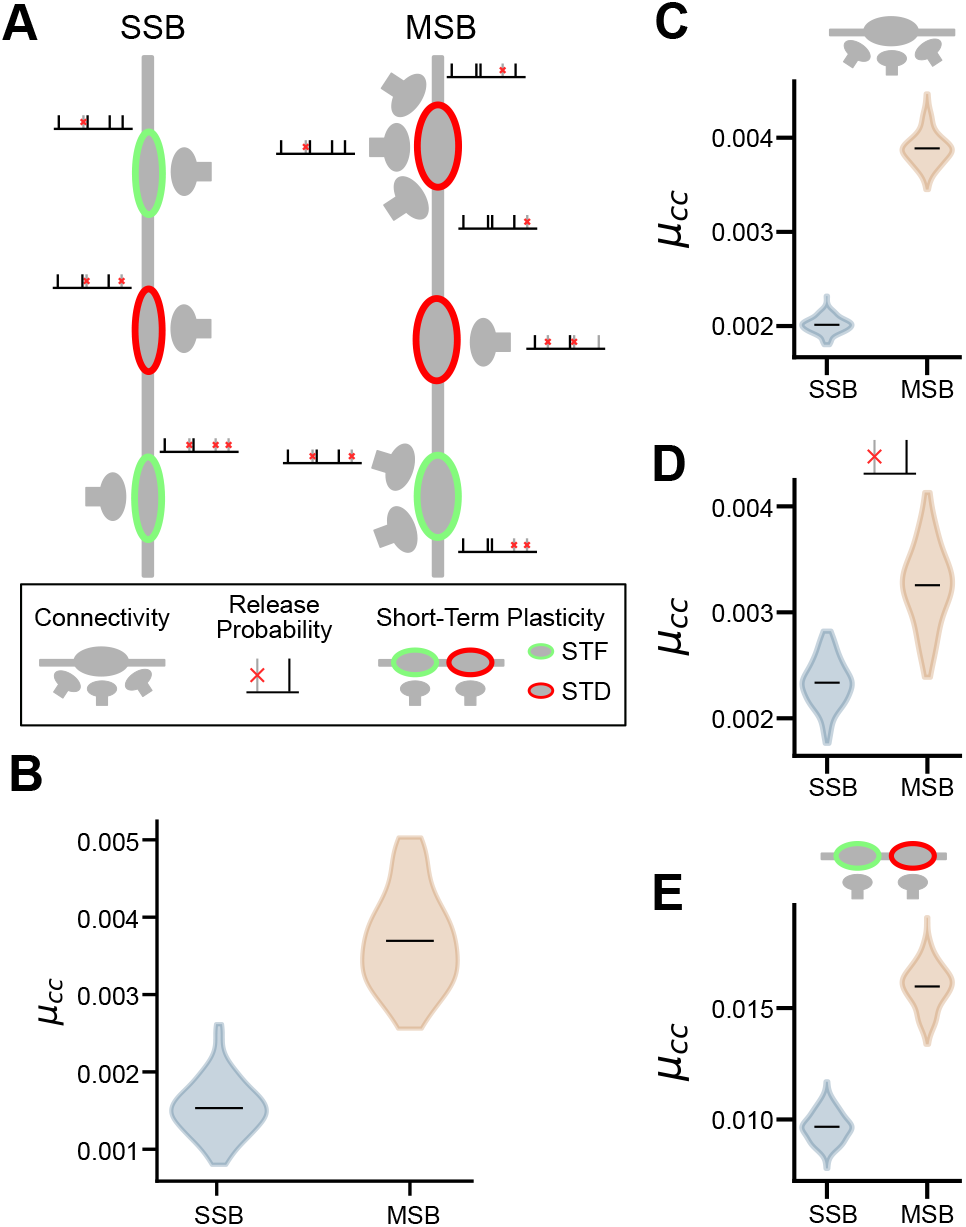
MSBs increase correlation in CA1 activity in computer simulations. (**A**) Schematic of the simulation. Neurons from CA3 have multiple boutons that synapse onto neurons in CA1. Each bouton undergoes either short-term facilitation (STF, green boutons) or short-term depression (STD, red boutons). Each bouton has a fixed release probability. In the SSB simulations, each bouton connects to a single CA1 neuron, whereas in the MSB simulations, each bouton might connect to multiple neurons in CA1. (**B**) Mean correlation coefficient (μ_cc_) between neurons in CA1 in the SSB and MSB simulations. (**C-E**) Same as B, but in setups with a single mechanism only: multiplicative connectivity (**C**), release probability (**D**), STP (**E**). (**B-E**) Grey area shows distribution of μ_cc_ across 100 independent simulation runs.

Next, we wanted to check if a single mechanism alone was responsible for the increase in correlation or if all three of them were contributing to the observed effect. We performed simulations in which only a single mechanism was active at a time and observed that all three of them had small positive effects on the correlation of networks with MSBs (Fig. 4C-E). Together, our results show that MSBs could indeed play a role in synchronizing CA1 activity and suggest multiple mechanisms could be involved in this synchronizing effect.

## Discussion

We show that a large proportion of synapses in the *stratum oriens* of the hippocampal CA1 subfield are MSBs, rather than the more typical SSBs. Previous EM reconstructions have described MSBs in a number of brain regions, including the hippocampus and cortex. In the *stratum radiatum* of the hippocampus, where these structures were originally described, only 24% of all excitatory synapses were found to be MSBs, of which 20% formed connections onto different dendrites (Sorra and Harris, 1993). Similarly, in the cortex, where a large number of synapses (^~^1700) have recently been annotated over a large 3D volume (0.3 mm^3^), 18% of all excitatory synapses were found to be MSBs (Kasthuri et al., 2015), in agreement with previous work (Knott et al., 2006). Although MSBs are generally not considered to be a feature of neuronal circuits in the brain, it is gradually becoming apparent that the textbook definition of the synapse does not necessarily apply to all brain regions (Bosch et al., 2015; Hara et al., 2011). Here, we show that for excitatory synapses in the *stratum oriens* (SO) of the hippocampal CA1 subfield, MSBs are present in roughly equal proportion to SSBs, suggesting they play a prominent role in circuit function.

The morphological properties of these MSBs reveal much about the possible role they play in the brain. From a network perspective, the MSBs described here are intriguing in that they contact spines from different dendrites that belong to different neurons. As a result, they act like synaptic transmission hubs, broadcasting signals across multiple CA1 neurons. Morphologically, MSBs are also unique in that the properties of synaptic contacts within an MSB are much less heterogeneous that between neighbouring SSBs. In consequence, MSBs are likely to transmit similar information across all AZs within a given bouton. Based on our mathematical models, we argue that these properties, when put together, all favour a role for MSBs in synchronizing activity across neurons in a network. CA1 neurons in the hippocampus show robust synchronous activity (Csicsvari et al., 2000), particularly across neighbouring cells (Takahashi and Sakurai, 2009), a feature that we suggest may emerge from the divergent connectivity pattern provided by MSBs.

Besides their role in synchrony, MSBs may also play an equally intriguing role in structural forms of synaptic plasticity. *In vitro* imaging has previously shown that spines formed onto MSBs are more dynamic than those onto SSBs (Reilly et al., 2011). This is particularly interesting when considering that *in vivo* measurements of dendritic spine turnover in the *stratum oriens* of the hippocampus is particularly high when compared to other brain regions such as the cortex (Attardo et al., 2015; Pfeiffer et al., 2018), a feature that may belie its role in the process of learning and memory. It is therefore tempting to speculate that the higher density of MSBs in the hippocampus provides the necessary substrate for this plasticity to take place. In essence, MSBs would act as the docking sites for spines to latch onto as they disappear and re-emerge during the acquisition, loss or retrieval of memories. In addition to this role, MSBs may also be well positioned to explain another peculiarity of synaptic plasticity in the hippocampus - the spread of LTP to neighbouring neurons (Bonhoeffer et al., 1989; Schuman and Madison, 1994). As previously proposed in an elegant review (Harris, 1995), the MSB could act as a node that spreads the potentiation of a given synaptic contact to all other synapses associated with it, through some presynaptic mechanism. The lower variance of synapses associated with a single MSB described here, further supports this notion. Although clearly these are early days, future experiments will need to uncover the role these unique structures play in the brain and the developmental processes that lead to their emergence.

## Acknowledgements

We would like to thank Winnie Wefelmeyer for useful feedback and comments on the manuscript. We would also like to thank Nicolas Bourg and Abbelight for the SAFeRedSTORM module and software, as well as Olympus for the IX3 microscope used for dSTORM. This research was funded in whole, or in part, by the Wellcome Trust (095589/Z/11/Z and 215508/Z/19/Z to JB). For the purpose of open access, the author has applied a CC BY public copyright licence to any Author Accepted Manuscript version arising from this submission. J.B.S and F.L. acknowledge the national infrastructure France BioImaging (ANR-10-INBS-04-01). This work was also supported by an ERC Starter Grant (282047) and a BBSRC project grant (BB/S000526/1) to JB, as well as an NC3Rs Training Fellowship to MR.

## Methods

### Animals

All animal procedures were approved by the local ethics committee and licensed under the UK Animals (Scientific Procedures) Act of 1986. Male and Female SV-129 mice were housed grouped in standard cages and provided with ad libitum food and water.

### Confocal imaging and dendrite tracing

Confocal imaging was performed using a Nikon A1 confocal microscope, equipped with a 40X water immersion objective (Olympus, Apo 40X WI λs DIC N3), and using NIS – Elements software. Images from 6 CA1 pyramidal cells sparsely labelled with Green Fluorescent Protein were obtained using high sampling resolution (pixel size 0.1 μm in X-Y and 0.3 μm or 0.8 μm in Z) and covering a large area of the basal dendritic tree (average area XY = 45551 μm2, average span Z = 45 μm).

Tracing was done using the Fiji Plugin Simple Neurite Tracer (Longair et al., 2011) and traces were smoothed and re-sampled at constant node inter-distance of 0.5 μm using MATLAB TREES Toolbox. Distances to the soma for each node were calculated using the Pvec_tree function of MATLAB TREES (Cuntz et al., 2010). Custom written functions in IgorPro 6.37 were written to calculate 3D Euclidean distances between each node and all the other nodes located in separate branches from the same dendritic tree. A threshold of 5 μm was used to determine the number of separate branches judged to be at a close enough distance to each node to contribute spines to multi-synaptic boutons. Nodes were ordered by distance to cell soma and average values across 10 distance bins comprising the same number of nodes is shown for each cell. A total distance of 8179 micrometres of dendrites was traced across 6 cells (average 1363 ± 158 micrometres per cell).

### Electron Microscopy

Two mice (post-natal day 22 and 100) were transcardially perfused with 20 ml of ice-cold saline solution followed by 200 ml of ice-cold fixative (2% PFA and 0.2 % glutaraldehyde mixture in 0.1 M phosphate buffer), followed by incubation overnight in fresh fixative at 4°C. Coronal vibratome sections (60 mm) were cut using a Leica VT1000S vibratome and further fixed in 1.5% potassium ferrocyanide: 2% osmium tetroxide in cacodylate buffer for 30 min at 4°C. Tissue was then thoroughly rinsed in distilled water and incubated in 1% aqueous thiocarbohydrazide for 4 min. After further rinsing, the samples were treated with 2% aqueous osmium tetroxide for 30 min, rinsed and en-bloc stained in 1% uranyl acetate for 2 h. To further enhance contrasts in the samples, one last treatment with Walton’s Lead was carried out for 30 min at 60°C, before proceeding to dehydration in an ethanol series and infiltration with Durcupan ACM resin (Sigma). After embedding and curing, tissue blocks were mounted on Gatan 3View aluminium pins using conductive glue (CircuitWorks Conductive Epoxy) and trimmed accordingly. Before imaging, samples were gold coated to increase electron conductivity. The specimens were then placed inside a Jeol field emission scanning electron microscope (JSM-7100F) equipped with a 3View 2XP system (Gatan). Section thickness was set at 40 nm (Z resolution). Samples were imaged at 2.5kV under high vacuum using a 2048×2048 scan rate, which gave a final pixel size of 4.4 nm.

Electron microscope images were registered and manually segmented using the ImageJ plugin TrakEM2 (Cardona et al., 2012). Extracted 3D structures were exported to the Blender software with the Neuromorph toolset (Jorstad et al., 2015), which was used to compute surface, volume and length measurements and render 3D reconstructions.

### Reconstruction of neurites

Stacks of SBFSEM images were concatenated and aligned using TrackEM plugin within the ImageJ environment (Chen et al., 2018). Portions of dendrites from proximal and distal regions of the stratum oriens were manually traced also using the TrackEM plugin. Only dendrites that could be followed for more than 5 um through the stack were traced. When a dendritic spine was formed onto a bouton, that bouton was traced, as well as any other dendritic spines that synapsed upon the same bouton. Extracted 3D structures were exported to the Blender software using the Neuromorph toolset to render 3D reconstructions (Jorstad et al., 2015) where surface area, volume, and length measurements were extracted. The P22 data set includes a subset of synapses included in a previous publication (Grillo et al., 2018), which was expanded for this study.

### Histology for dSTORM

Mice (22 days old) were anaesthetized with an overdose of sodium pentobarbital and transcardially perfused with 30 mL of ice-cold saline solution followed by 40 mL of 4% (w/v) PFA (pH 7.3). The brains were carefully removed and put in 4% (w/v) PFA solution overnight at 4°C. Brains were incubated in 15% sucrose then in 30% sucrose, both overnight at 4°C. They were embedded in blocks of Gelatine/Sucrose and froze in isopentane (−60°C) and then stored at −80°C until cryo-sectioning. Coronal slice of 40μm thick were obtained with a cryostat (Leica). Slices were kept in anti-freeze PBS at −20°C. On the day of staining, slices were incubated with warm PBS 4 times 5 minutes each, to remove the gelatine/sucrose blocks. A 50mM NH4Cl solution was used to quench the remaining PFA. An antigen retrieval step was performed by incubating brain slices in Citrate buffer (10mM sodium citrate, 0.05% Tween20, pH6) at 85°C for 25 minutes, followed by 3 PBS washes. For permeabilization, sections were incubated 4 times in a 0.25% Triton X-100 solution for 15 minutes each at room temperature (RT) on a shaker. Brain slices were incubated in a Blocking Buffer (5% BSA, 0.25% Triton X100, 10% Goat Serum) for 2 hours at RT on a shaker. This was followed by an overnight incubation with primary antibodies diluted in an Antibody Buffer (1% BSA, 0.25% Triston X-100, 5% Goat serum). On the following morning, slices underwent four 15 minutes PBS-Triton X-100 (0.1%) washes, followed by a 2-hour incubation at room temperature with secondary antibodies diluted in the Antibody Buffer. The secondary antibody solution was washed off as before and slices were mounted onto 1.5H 25mm glass coverslip (Marienfeld). Slices were allowed to dry on the coverslips, melted 2% agarose was used to immobilize the tissue, and coverslip were kept in PBS at 4°C until imaging.

VGlut was labeled using an anti-VGlut1 (Rabbit, Synaptic Systems – 135 303) diluted at 1/500 and an anti-Rabbit coupled to CF680 (Goat, Biotium – 20818) diluted at 1/1000.

Bassoon was labeled using an anti-Bassoon (Mouse, Abcam – ab82958) diluted at 1/500 and an anti-Mouse coupled to AF647 (Goat, Invitrogen – A32728) diluted at 1/1000.

### Multicolor 3D-dSTORM imaging

For dSTORM, coverslips were placed in an imaging chamber (AttoFluor Cell Chamber, Invitrogen). The imaging chamber was filled with STORM Buffer and sealed using another glass coverslip. Final dSTORM imaging buffers were composed of oxygen scavengers (100 μg/ml Glucose oxidase (Sigma-Aldrich G2133) and 4 μg/ml Catalase (Sigma-Aldrich C100)) and reductive agent (100mM β-Mercaptoethylamine-HCl, Sigma-Aldrich M6500). Oxygen scavengers stock solution containing 20mM Tris–HCl pH 7.2, 4mM TCEP, 25mM KCl, 50% Glycerol and 1mg/ml Glucose oxidase (Sigma-Aldrich G2133) and 42 μg/ml Catalase (Sigma-Aldrich C100) and was stored at −20°C. Reducing agent stock solution containing 1M β-Mercaptoethylamine-HCl (Sigma-Aldrich M6500) in deionized water and pH adjusted at pH8 with NaOH, was stored at −20°C. Dilution STORM Buffer containing 100mg/mL Glucose (Sigma-Aldrich) and 10% Glycerol (Fisher Scientific) in deionized water, was stored at 4°C.

Multicolor dSTORM imaging was performed using a spectral-demixing SAFeRedSTORM module (Abbelight, France) mounted on an Olympus IX3 equipped with an oil-immersion objective (100× 1.5NA oil immersion, Olympus). The two fluorophores, AF647 and CF680 were excited with a single wavelength using a fibber-coupled 642nm laser (450mW Errol, France). Because the two fluorophores have a small shift in their emission spectrum, a long-pass dichroic beam splitter (700nm; Chroma Technology) was used to split the emission light on two cameras (ORCA-fusion sCMOS camera (Hamamatsu)). A photon ratio was calculated for each detection to assign the detection to one of the fluorophores (Lampe et al., 2012; Testa et al., 2010). 3D-dSTORM imaging was performed by using cylindrical lenses placed before each camera.

Image acquisition and control of microscope hardware were driven by Abbelight’s NEO software. Region of interest (ROI) of 512 × 512 pixels (pixel size = 97nm) within the stratum oriens were identified. Depth of imaging was limited to the first ten micrometres of the slice due to the working distance of the objective and to allow a proper focus stabilization using the Olympus’s ZDC system. Image stacks contained 60,000 frames. Cross-correlation was used to correct for lateral drifts. N 3D calibration was performed using fluorescent beads (tetraspeck beads, invitrogen) to measure the x,y deformation obtained with the 2 cylindrical lenses on both cameras. With this information we calculated the z position of individual localizations with a precision of ^~^1μm. Together with a gaussian fit to single emitter events in the x-y plane (Maximum Likelihood Estimation, MLE, using the NEO_analysis software) we were able to obtain a the 3D spatial coordinates of single molecules with high resolution. A first transformation (cross-correlation) was applied to precisely realign detection events from both cameras. Finally, for each detection, a ratio of intensity between each camera was calculated (I1 / (I1+I2)) in order to reassign the detection to one of the two fluorophores. Superresolution images (.tiff) with a pixel size of 10nm and localization files (.csv) containing the 3D spatial coordinates of each demixed detection (detection of each fluorophores) were generated.

### Multicolor 3D-dSTORM analysis

Point Clouds Analyst (PoCA) (Levet, 2021), a continuation of SR-Tesseler (Levet et al., 2015) and Coloc-Tesseler (Levet et al., 2019), was used to identify Single Synaptic and Multi Synaptic Boutons and to quantify VGlut (color 1) and Bassoon (color 2) clustering from localized molecule 3D coordinates. For each color, a 3D Voronoi diagram was computed by creating polyhedrons of various sizes centered on the localized molecules. Automatic segmentation allowed detecting clusters for each protein (VGlut cluster to identify excitatory presynaptic boutons, and Bassoon clusters to identify Active Zones) by using a density threshold ^2^ for each colour (δi1 and δi2, respectively) based on the mean density of molecules for the whole dataset, such that δi1 ≥ 4δd1 and δi2 ≥ 2δd2 for color 1 (VGlut) and color 2 (Bassoon), respectively. We only selected clusters with a minimum number of localizations, set to 200 and 50 localizations for color 1 and color 2, respectively. The colocalization between VGlut and Bassoon was computed from the overlapping clusters. Because of the stringent segmentation performed on color 1 (VGlut), which was needed due to the high density of synapses in brain slices, a significant amount of color 2 clusters (Bassoon) did not colocalize with color 1 clusters (Supplemental Figure 3). To avoid colocalization issues, we manually picked 480 synapses of those identified automatically, where there was no sign of miscolocalization (see Supplemental Figure 3), half of which were MSBs. Synapses were picked on multiples images from different brain slices from 3 mice. From these synapses, we extracted cluster volumes and localization densities for each color. Variances in cluster volume and density of multiple Bassoon clusters colocalizing with a single VGlut cluster (MSB) were measured and compared to randomized single Bassoon clusters which colocalized with a single VGlut cluster (SSB).

### Data Analysis and Statistics

Data analysis was performed, and graphical visualisation made, using R 3.4.1 (R Foundation for Statistical Computing; http://www.r-project.org/) and RStudio 1.2.5042 (https://rstudio.com).

Single variable data distributions were visualised using kernel density estimation plots. This non-parametric probability density function provided a more effective way to view the distribution of a variable.

Pooled synapse density and morphological measurements were visualised using violin or box plots. Box-and-whisker plots indicate the median (thick horizontal line), the 25th–75th percentiles (box) and the 10th–90th percentiles (black whiskers); circles within and to the right of the box represent mean and individual values, respectively. Violin plots are comparable to box plots, but give a representation of the kernel probability density of the data. Grey or black lines that extend from the centre to the plot perimeter indicate the median of the separated MSB and SSB data. The box inside the violin represents the median and interquartile range of the combined MSB and SSB data.

Comparisons between synaptic densities and synapse morphologies were visualised graphically using estimation plots of effect sizes, and their uncertainty assessed using confidence intervals (Ho et al., 2019). This approach was preferred over traditional null hypothesis significance testing because of the associated limitations (Goodman, 2008). Separation between the confidence interval and 0 was considered an effect. Confidence intervals were adjusted for multiple comparisons by an extension of the Ryan–Holm step-down Bonferroni procedure. Confidence intervals were adjusted for 4 comparisons for measures of synaptic density, whereas 12 comparisons were corrected for measures of synapse strengths. Differences between SSB and MSB synaptic strength variances were tested using Cliff’s delta and shown using Gardner-Altman estimation plots. 5000 bootstrap samples were taken; the confidence interval was bias-corrected and accelerated. For each permutation P value, 5000 reshuffles of the control and test labels were performed.

Hierarchical cluster analysis was performed with the ward minimum variance method using the nbclust package in R that aggregates 30 indices (Charrad et al., 2014).

Correlations were tested using Spearman’s rank-order correlation. P and r values are presented as equalities (two significant figures).

### Computational Model

All simulations were performed using the neural network simulator NEST 2.20.0 (Fardet et al., 2020) and the code will be released on GitHub after publication.

#### Neuron model

Neurons from CA1 are current based leaky integrate-and-fire (LIF) with exponential postsynaptic currents. The sub-threshold membrane potential V of a neuron obeys the following equation:

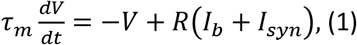

where *τ_m_*= 60 ms is the membrane time constant and R = 1GOhm is the input resistance. The background input *I_b_* is modeled as a Gaussian white noise current with mean *μ_b_* = 18 pA and standard deviation *σ_b_* = 20 pA. The input current from presynaptic neurons *I_syn_* is modeled as the sum of input currents from all presynaptic neurons. Every time the membrane potential reaches a threshold value *V_th_* = 20mV, the neuron emits a spike. Following a spike, the membrane potential is reset to *V_reset_* = 10mV, and remains there for a refractory period *t_ref_* = 2 ms. In simulations without short-term plasticity (STP), the input current from a presynaptic neuron consists of their spike train filtered with an exponential with maximum amplitude A = 200 pA and time constant *τ_syn_* = 1.5 ms. Neurons from CA3/2 are modeled as independent spike trains with Poisson statistics and rate r = 10 Hz.

#### Short-term plasticity

Where synapses are plastic, we use the STP model implemented on NEST 2.20.0 (Fardet et al., 2020), according to (Tsodyks et al., 2000). The total synaptic input to a postsynaptic neuron *i* is given by:

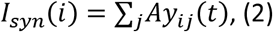

where *A* is the absolute synaptic weight, and *y_ij_* determines the effective contribution of the postsynaptic current (PSC) from neuron *j* to the input current to neuron *i*. It evolves according to the system of equations:

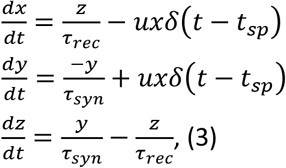

where *x, y* and *z* are the fraction of synaptic resources in the recovered, active and inactive states respectively, *t_sp_* denotes the timing of a presynaptic spike, *τ_syn_* is the decay time constant of PSCs and *τ_rec_* is the recovery time constant for depression. The variable *u* describes the effective use of synaptic resources by each presynaptic spike, and it evolves according to:

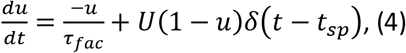

where *τ_fac_* is the time constant for facilitation. The parameters used were extracted from (Tsodyks et al., 1998). For synapses undergoing facilitation, A = 1540 pA, U = 0.03, *τ_rec_* = 130 ms, *τ_fac_* = 530 ms. For synapses undergoing depression, A = 250 pA, U = 0.5, *τ_rec_* = 800 ms, *τ_fac_* = 0 ms.

#### Correlation coefficient

We calculate the Pearson’s correlation coefficient between spike trains of every pair of postsynaptic neurons. All simulations are run for 51 s. Spike trains are created from the last 50 s of simulation, with bins of 20 ms. Each independent simulation run generates a single mean value of correlation coefficient *μ_cc_*, across all pairs of postsynaptic neurons. Figures show distribution of *μ_cc_* across 100 independent simulation runs.

#### Full simulation

We simulate two networks representing CA1 and CA3/2, with 100 excitatory neurons each. CA3/2 neurons are modeled as independent spike trains with Poisson statistics and rate r = 10 Hz. CA1 LIF neurons receive a background input *I_b_* and synaptic inputs *I_syn_* from CA3/2 neurons. Each neuron from CA3/2 has 5 boutons. Each bouton may have one single or multiple active zones, as described in the following, and each active zone always connects to a singe postsynaptic CA1 neuron. *Multiplicative*: in the SSB network, each bouton has only one active zone which connects to one randomly chosen postsynaptic CA1 neuron. In the MSB network, each bouton may have a number *α* of active zones, where *α* is randomly selected from the interval {*α* ∈ *Z*: 1 ≤ *α* ≤ 5} with probabilities extracted from Figure 1C. Each active zone connects to one randomly chosen postsynaptic CA1 neuron. *Release probability*: Each bouton *i* has a release probability *p_i_*, which is randomly chosen from a Gamma distribution with parameters shape *k* = 2 and scale *θ* = 0.15, and with values bounded to a maximum of 1 (Figure SA). The spike trains at different active zones within the same bouton *i* are different realizations of sampling from the same presynaptic spike train with the same probability *p_i_*. *STP*: all connections are subject to STP. Each bouton is either undergoing short-term facilitation or short-term depression. The short-term plasticity state of each bouton applies to all active zones within that bouton.

#### Multiplicative only simulation

These simulations are the same as the full simulation, except that: (i) release probability *p_i_* = 1 for all boutons; and (ii) there is no STP.

#### Release probability only simulation

These simulations are the same as the full simulation, except that: (i) each presynaptic neuron from CA3/2 has either 5 boutons with a single active zone each (in the SSB scenario) or one bouton with 5 actives zones (in the MSB scenario). In both SSB and MSB scenarios, each active zone connects to one randomly chosen CA1 postsynaptic neuron; and (ii) there is no STP.

#### STP only simulation

These simulations are the same as the full simulation, except that: (i) each presynaptic neuron from CA3/2 has either 5 boutons with a single active zone each (in the SSB scenario) or one bouton with 5 actives zones (in the MSB scenario). In both SSB and MSB scenarios, each active zone connects to one randomly chosen CA1 postsynaptic neuron; and (ii) release probability *p_i_* = 1 for all boutons.

---

**Figure S1.**
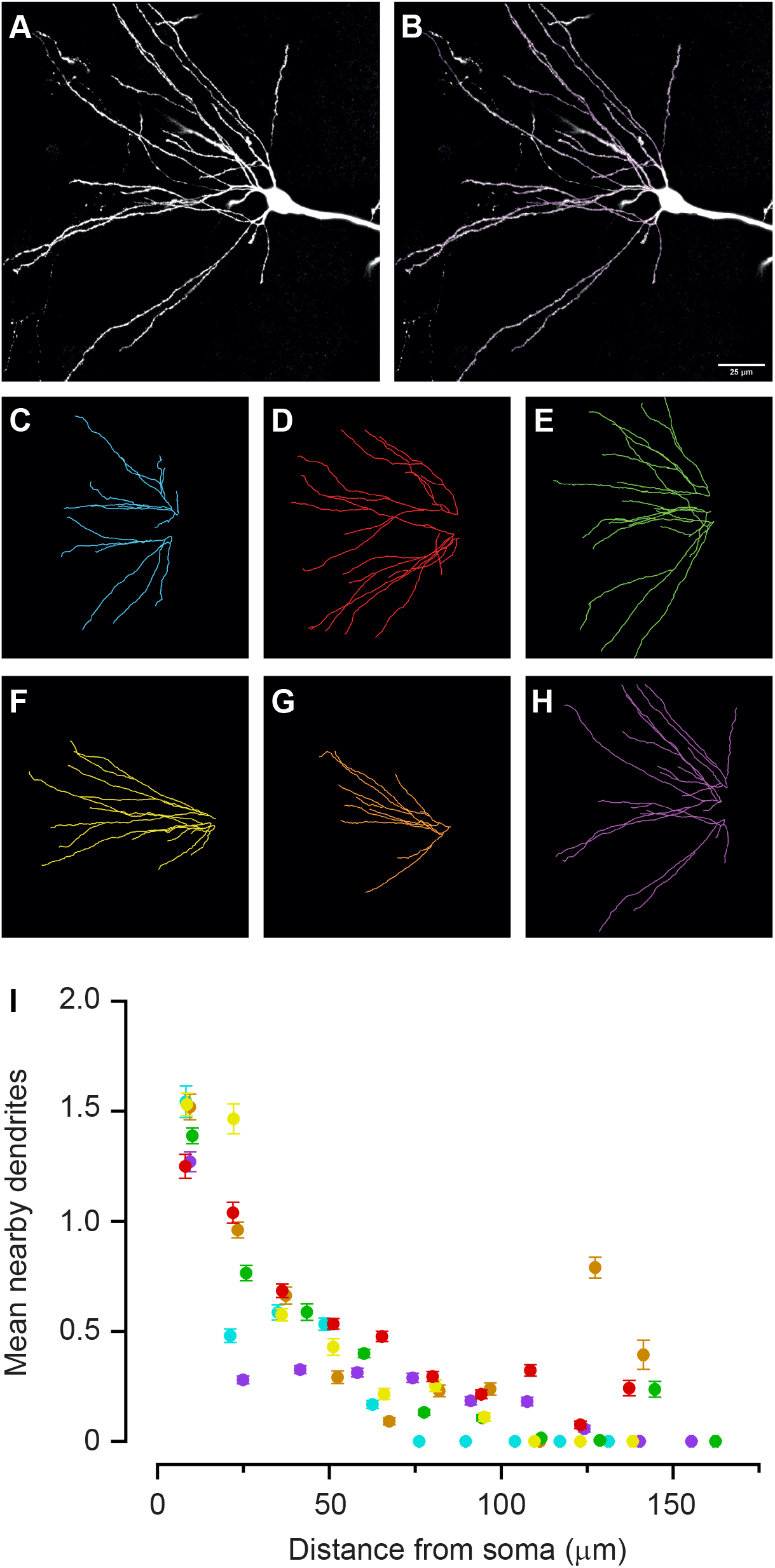
The basal dendrites of a single CA1 hippocampal neuron are unlikely to contribute multiple synapses to MSBs. (**A**) Example image of the basal dendrites of a CA1 pyramidal neuron transfected with EGFP using in utero electroporation. (**B**) Tracing of the basal dendrites of the neuron shown in A. (**C-H**) Maximal intensity projections of the tracings of the basal dendrites of 6 different CA1 pyramidal neurons used in this analysis. (**I**) A plot of the number of nearby dendrites with respect to a reference dendrite from the same cell, as function of distance from the soma. A 5 μm radius sphere (roughly 2 spine lengths) was moved along a reference dendrite and used as a way of assessing the proximity of dendrites from the same cell that could contribute to an MSB. A value of 1 indicates that a single other dendrite fell within the sphere and was therefore close enough to potentially contribute a spine to an MSB. The graph clearly shows that beyond 50 μm distance from the soma, the number of potential contacts is very low and cannot explain that large numbers of MSBs described here.

**Figure S2.**
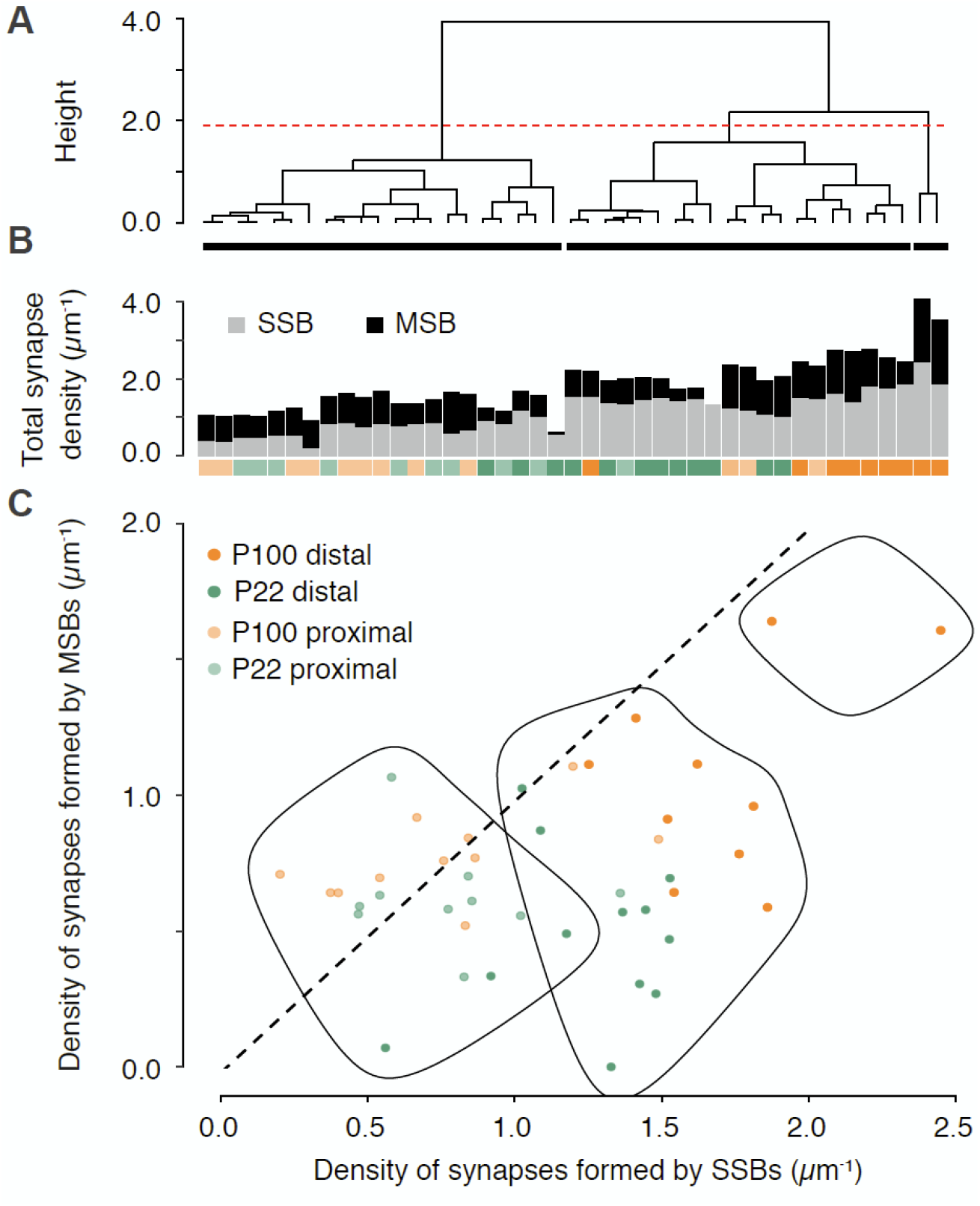
Hierarchical clustering of synapses along basal dendrites show importance of synapse position. (**A – C**) Hierarchical cluster analysis using the ward minimum variance method was used to determine if dendrites could be separated by their densities of single synaptic and multi synaptic boutons. By using the nbclust package in R that aggregates 30 indices (Charrad et al. 2012) the data was most appropriately categorised into 3 groups (red dotted line); dendrites that form roughly equal numbers of synapses onto MSBs and SSBs, dendrites that form more SSBs than MSBs and 2 dendrites which have a high density of both MSBs and SSBs. Without information on their region or age, dendrites quite accurately clustered according to whether they are distally or proximally located relative to CA1 somata, with distal dendrites forming equal contacts with MSBs and SSBs, whilst proximal dendrites tended to form spines onto SSBs.

**Figure S3.**
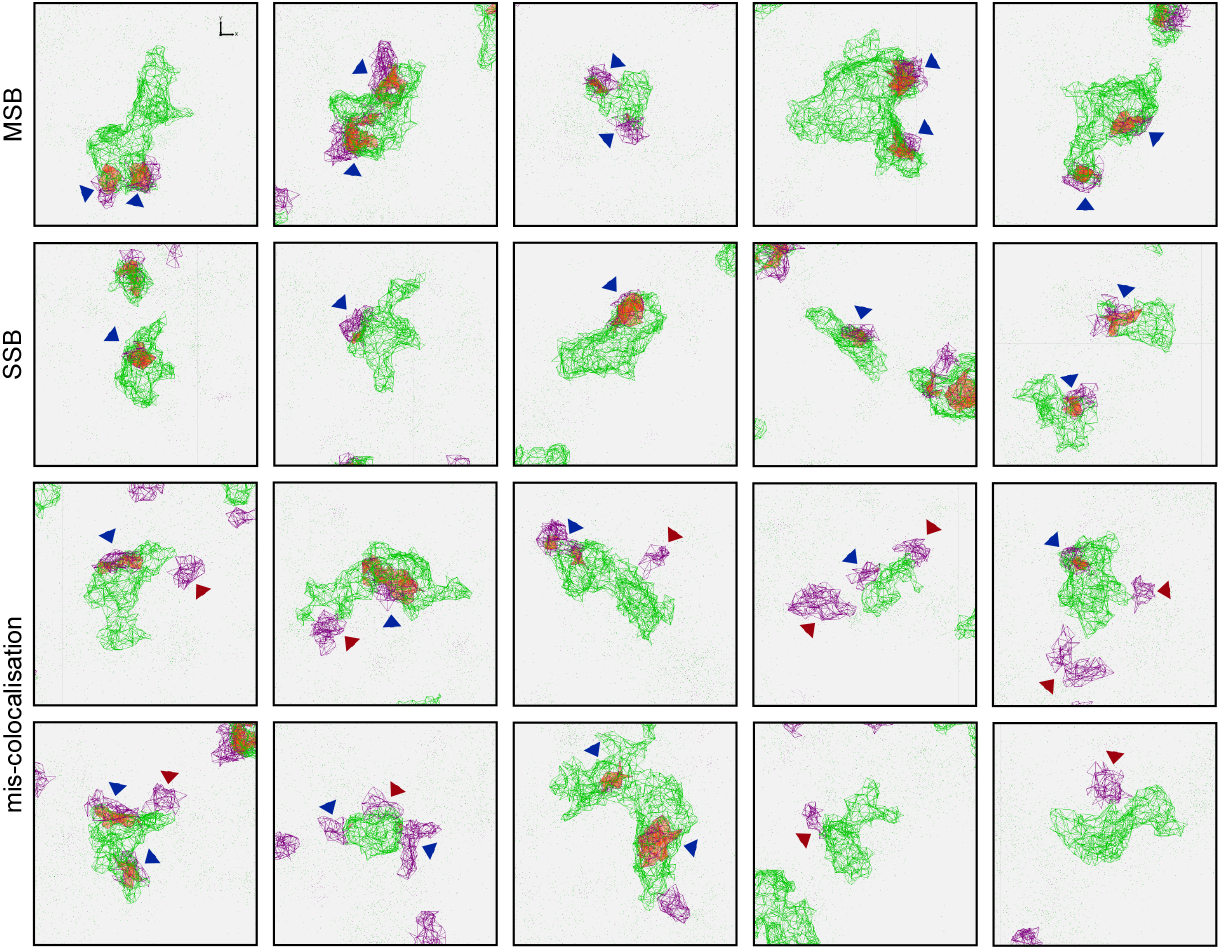
Challenges associated with co-localisation of segmented clusters and their assignment into MSBs and SSBs. Multiple examples of identified MSBs (first row) and SSBs (second row), showing correct automated colocalisations (blue arrow heads). Third and fourth rows show putative colocalization errors from Bassoon puncta lying close, but outside a VGlut cluster (red arrow heads). This could result in an MSB being incorrectly assigned as an SSB (third row), in an MSB with an incorrect number of AZs (fourth row, examples 1 to 3), or in a bouton without any AZs (fourth row, examples 4 and 5). We took a conservative approach that only picked segmented clusters showing direct colocalisation (blue arrows). This approach will underestimate MSBs but avoids any false positive detections.

**Figure S4.**
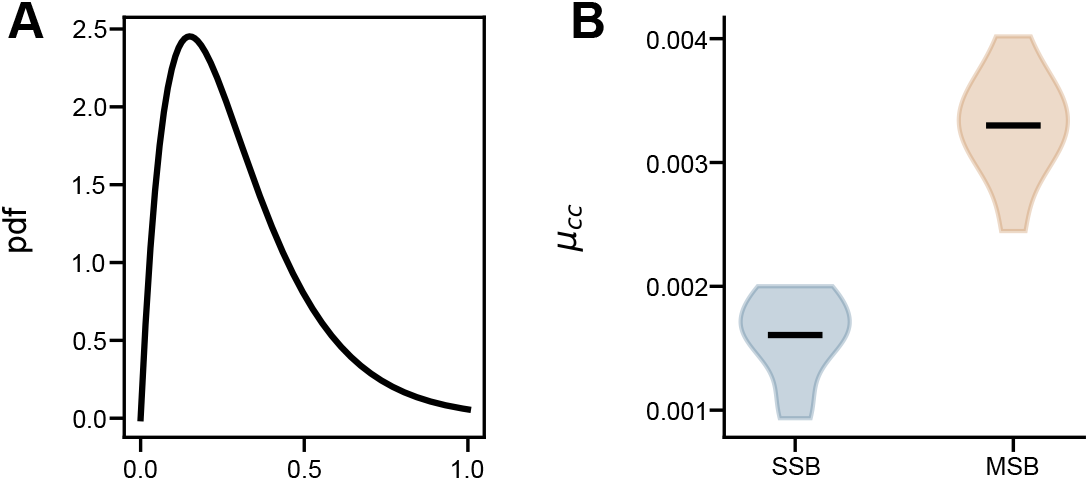
MSBs increase the correlation of CA1 neurons when the STP of a bouton is linked to its Pr. (**A**) Gamma distribution from which values of release probabilities are drawn. (**B**) Same as Figure 4B, but with non-independent release probability and short-term plasticity. The main model (Fig. 4) implicitly assumes that the short-term plasticity state of each bouton is independent of their release probability. As an alternative, here we simulate a scenario in which they are not independent. For each bouton, we draw a value of release probability *p_i_*, which decides its short-term plasticity state: boutons are depressing if *p_i_* ≥ 0.5 and facilitating otherwise.

